# Whole genome survey of big cats (Genus: *Panthera*) identifies novel microsatellites of utility in conservation genetic study

**DOI:** 10.1101/2020.07.08.193318

**Authors:** Jee Yun Hyun, Puneet Pandey, Kyung Seok Kim, Alvin Chon, Daecheol Jeong, Jong Bhak, Mi-Hyun Yoo, Hey-Kyung Song, Randeep Singh, Mi-Sook Min, Surendra Prakash Goyal, Damdingiin Bayarkhagva, Hang Lee

## Abstract

Big cats (Genus: *Panthera*) are among the most threatened mammal groups of the world, owing to illegal transnational trade. Conservation genetic studies and effective curbs on poaching are important for the conservation of these charismatic apex predators. A limited number of microsatellite markers exists for *Panthera* species and researchers often cross-amplify domestic cat microsatellites to study these species. We conducted data mining of seven *Panthera* genome sequences to discover microsatellites for conservation genetic studies of four threatened big cat species. A total of 32 polymorphic microsatellite loci were identified in silico and tested with 99 big cat individuals and 7 Eurasian lynx. The developed markers were polymorphic in most of the tested species. We propose a set of 15 novel microsatellite markers for use in conservation genetics and wildlife forensic investigations of big cat species. Cumulatively, these markers have a high discriminatory power of one in a million for unrelated individuals and one in a thousand for siblings. Similar PCR conditions of these markers increase the prospects of achieving efficient multiplex PCR assays. This study is a pioneering attempt to synthesise genome wide microsatellite markers for big cats.

## Introduction

The genus *Panthera* includes five hyper carnivorous apex predator species that are typically referred to as big cats ^1–3^. These are the tiger (*Panthera tigris*), leopard (*Panthera pardus*), lion (*Panthera leo*), snow leopard (*Panthera uncia*), and jaguar (*Panthera onca*). Big cats have great conservation value. They play a significant role in ensuring proper ecosystem function through top-down regulation ^4^. Being charismatic, big cats helps in mobilising mass audiences and funds for the conservation cause ^5^. Also, big cats are species of great cultural and historical significance with references found in artwork, folk tales, and old sayings throughout their distribution ^6,7^. Nevertheless, a rampant decline in their wild populations has been observed in the recent past, mainly due to excessive hunting (including the prey species) and overexploitation of habitat resources. From 1970 onward, several measures have been undertaken globally to fight the cause of this falloff. These include a (1) hunting and trade ban, (2) periodic population census, (3) regional and international cooperation to initiate activities for habitat restoration and reintroductions, and (4) community sensitisation campaigns to mitigate conflict with humans ^8–12^. However, the success of such measures has been limited as these species continue to be listed among the IUCN (International Union for Conservation of Nature) endangered species ^13–17^.

Illegal Wildlife Trade (IWT) of big cats is a highly lucrative and unlawful transnational commercial activity that is worth millions of dollars annually ^18^. Though there is a moratorium on hunting and trade of big cat species (except African lion), they are poached in range countries, and their parts and products smuggled to China and Southeast Asia to meet the demands of businesses engaged in the manufacturing of traditional medicines, home decors, and ceremonial clothing ^19,20^. Between 2001 to 2010, covert investigations have found 493 big cat parts in the markets of Thailand and Myanmar ^21^. Similarly, 43 snow leopard seizures with at least 100 individuals have been reported in Chinese media between 2000 to 2013 ^22^. In the past decade, an estimated 6,000 African lion (only big cat species listed in Appendix II of CITES – Convention on International Trade of Endangered Species of Wild flora and fauna) skeletons have been moved legally (using CITES permit) to Southeast Asia and marketed as an alternative to tiger bones ^23^. As per the TRAFFIC report published in 2016, 810 tiger seizures have been made by law enforcement agencies between 2002 to 2013, accounting for more than 1,700 individuals ^24^. Poaching-driven regional tiger extinctions have occurred in India, Cambodia, Vietnam, Thailand, Korea, and other Asian countries in the past two decades ^25–28^. Regulations (i.e. national laws, international treaties, and conventions) have failed to curb the illegal trade of big cats as this illicit trade is a complex, fast-evolving and a heterogeneous transnational issue involving multiple trading partners/middlemen. Traded articles mostly lack morphological features to ascertain the species, reducing the ability to track their origins reliably.

Incremental adoption of genetic tools and techniques for wildlife conservation and management have been observed globally in the past 25 years mainly due to the development of the robust protocols for DNA extraction and PCR (Polymerase chain reaction) ^29–32^. DNA tools are now increasingly employed for establishing species-level identity ^33,34^, resolving taxonomic ambiguities^6,35,36^, wildlife conflict mitigation^37,38^, and more recently, establishing the source of origin ^39–41^. Microsatellites or short tandem repeats (STR) are neutral, co-dominantly inherited, widely distributed, hypervariable, short repetitive nuclear DNA units that have been regarded as the best candidate to develop a genetic signature of the individual (DNA fingerprint), population, and subspecies. Multiplex STR systems to undertake geographic assignments of confiscations have been proposed for tigers, leopards, elephants, rhinos and many other endangered species ^39,41–45^. However, except for rhinos and elephants, microsatellite-based applications have failed to achieve global consensus in wildlife offense investigation. Efficient and simple protocols with established utilities in wildlife forensics across the range and species of rhinos and elephants have convinced wildlife managers and law enforcement agencies to adopt DNA methods for seizure investigations.

Tiger, leopard, lion, and snow leopard are the four most commercially exploited (by poaching and illegal trade) *Panthera* species. Their conservation demands stringent law enforcement. Here, we report the development of novel microsatellite markers for genus *Panthera* by mining the genome sequences of four (tiger, leopard, lion, and snow leopard) most exploited big cat species. This study is a part of an ongoing India-Korea-Russia collaborative initiative to develop and test microsatellite based multiplex PCR panels of the pantherine species for genetic identification of the whole genus *Panthera*.

## Results

### Abundance and distribution of STR in genomes of big cat species

We analysed the whole genome sequences of seven big cat individuals and found a total of 80,474,871 variant sites. These include SNVs (single nucleotide variants), indels, and microsatellites. Potential target variants were mined within these variant sites following the protocols described in the materials and methods section. Some of these variants were consistently polymorphic across all genomes, whereas some had limited polymorphism. Due to a large number of potential target variant candidates, we selected only those that were at least polymorphic in 5 of the 7 big cat genomes. Altogether, there were 8,947 such potential target variants. Of these, 6,283 were found to be located on unique sites in the genome (unique target variant, UTV). We found 2,614 UTVs in all seven genomes, and these were finally processed for microsatellite screening using the program MSDB ^46^.

In big cat genomes, the dinucleotide microsatellite repeats were most abundant (45.4%), followed by mononucleotides (32.7%) and tetranucleotides (11.1%) (Fig. 2). The trinucleotides (8.6%), pentanucleotides (1.9%), and hexanucleotides (0.3%) were found in less abundance (Fig. 2). Relative abundance (mean number of STRs per Mb of genome analysed) was found to be the highest for Bengal tiger (357.3 STR/Mb) followed by white tiger (355.2 STR/Mb), Amur leopard (336.2 STR/Mb), Amur tiger (316.9 STR/Mb), white lion (312.3 STR/Mb), lion (310.7 STR/Mb), and snow leopard (304.4 STR/Mb).

**Figure 1:**
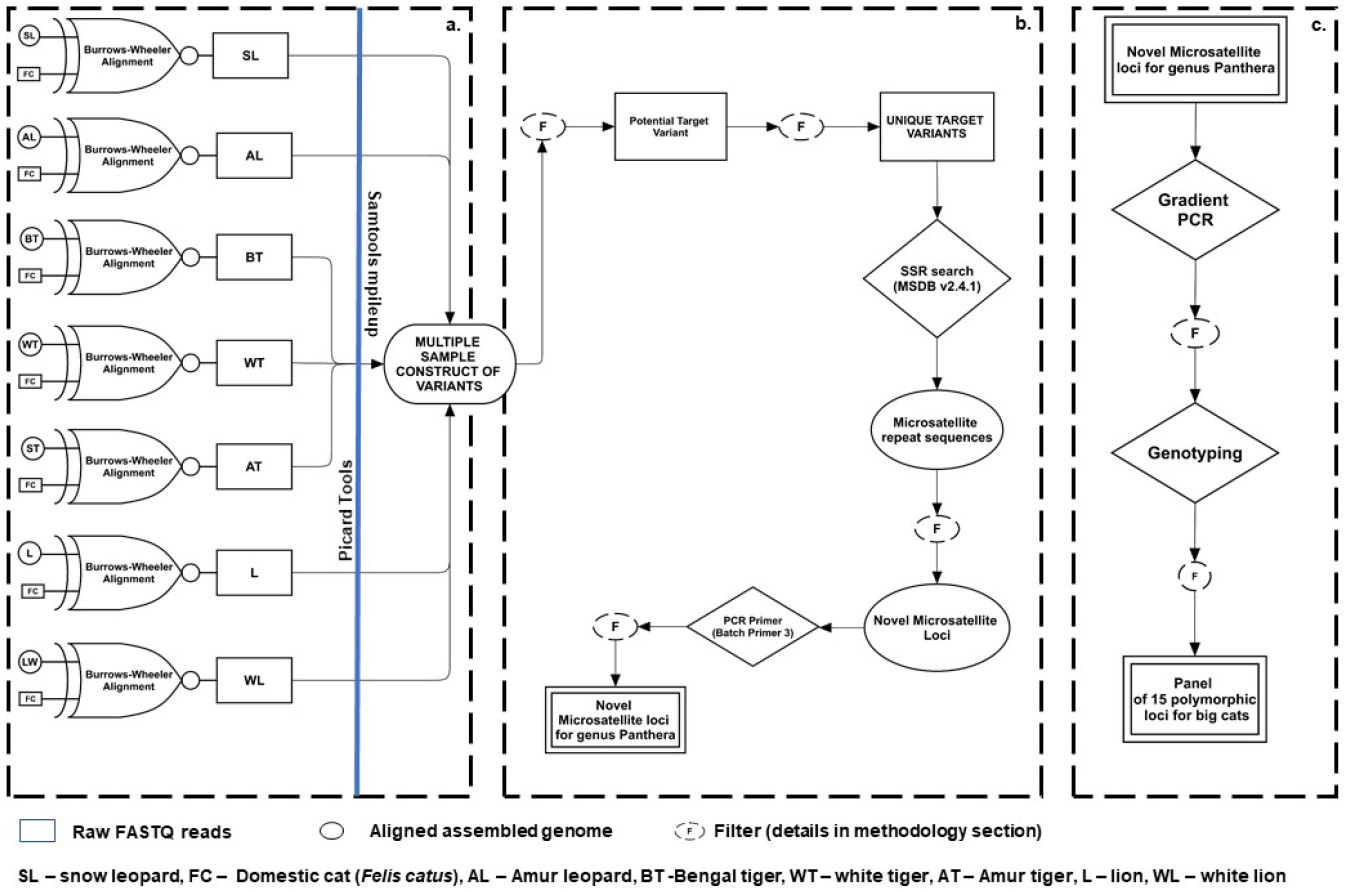
Schematic illustration of the STR marker development pipeline used in this study – (a) Genome alignment, (b) Microsatellite repeat search and PCR primer designing, and (c) Amplification and polymorphism evaluation of novel microsatellites.

**Figure 2:**
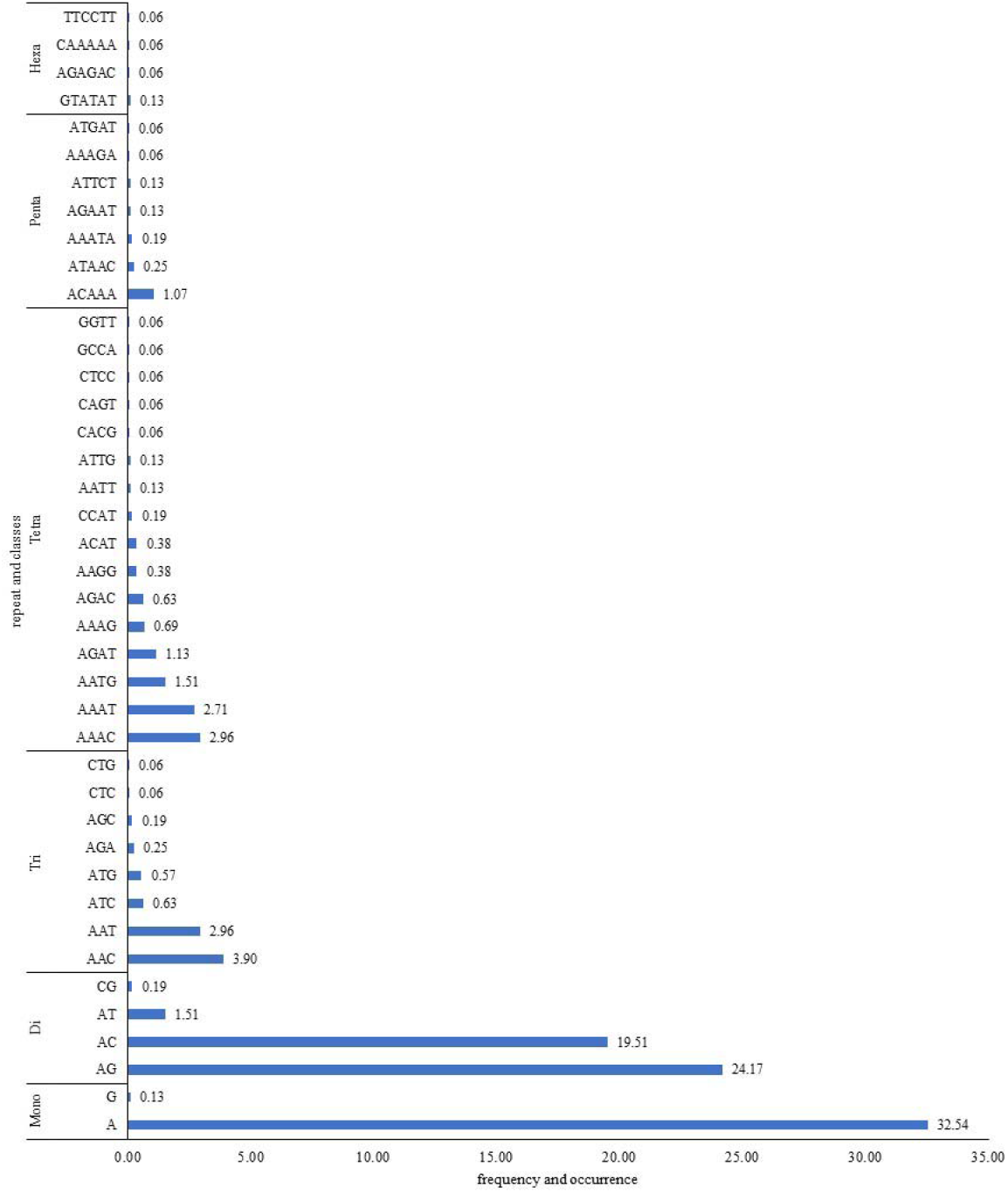
Frequency of occurrence of different STR repeat type classes across the *Panthera* genomes

Among all the mononucleotide repeats, (A)n was the most abundant (99.6%), while (C)n was comparatively scarce. In the dinucleotide repeat category, (AG)n and (AC)n were the two most frequent (96.3%) microsatellite motifs. Almost 80% of the trinucleotide types were (AAC)n, and (AAT)n in the *Panthera* genomes. Nearly half of the tetranucleotides were (AAAT)n and (AAAC)n. Among pentanucleotides, (ACAAA)n was the most abundant (56.7%). Hexanucleotides were the least among all types of microsatellites screened. The three most abundant microsatellite classes were (A)n, (AG)n, and (AC)n. Together they comprise 76.2% of the all forty-one microsatellite classes identified.

### Development of microsatellite markers for genus Panthera

Program batch primer 3 was used to design PCR primers ^47^. About 4% of the UTVs were found suitable for primer design (i.e. sufficient flanking sequences and not single-copy sequences). These include 176 dinucleotides, 39 trinucleotides, 45 tetranucleotides, 11 pentanucleotides, and 3 hexanucleotides. The designed primer pairs for these loci were further screened based on GC content and the presence of secondary structures. Finally, primer pairs for 41 loci were shortlisted for oligonucleotide synthesis. PCR was subsequently attempted with the synthesised primer pairs with four DNA samples, one each of the tiger, leopard, lion, and snow leopard. Thirty-two microsatellite loci (Table 1) showed clear amplification in the expected size range and were considered further. The forward primers of these loci were fluorescently labelled with one of the four dyes – 6FAM, VIC, NED, and PET. These labelled microsatellites were then used to genotype samples of tiger, leopard, lion, snow leopard, and lynx.

**Table 1:**
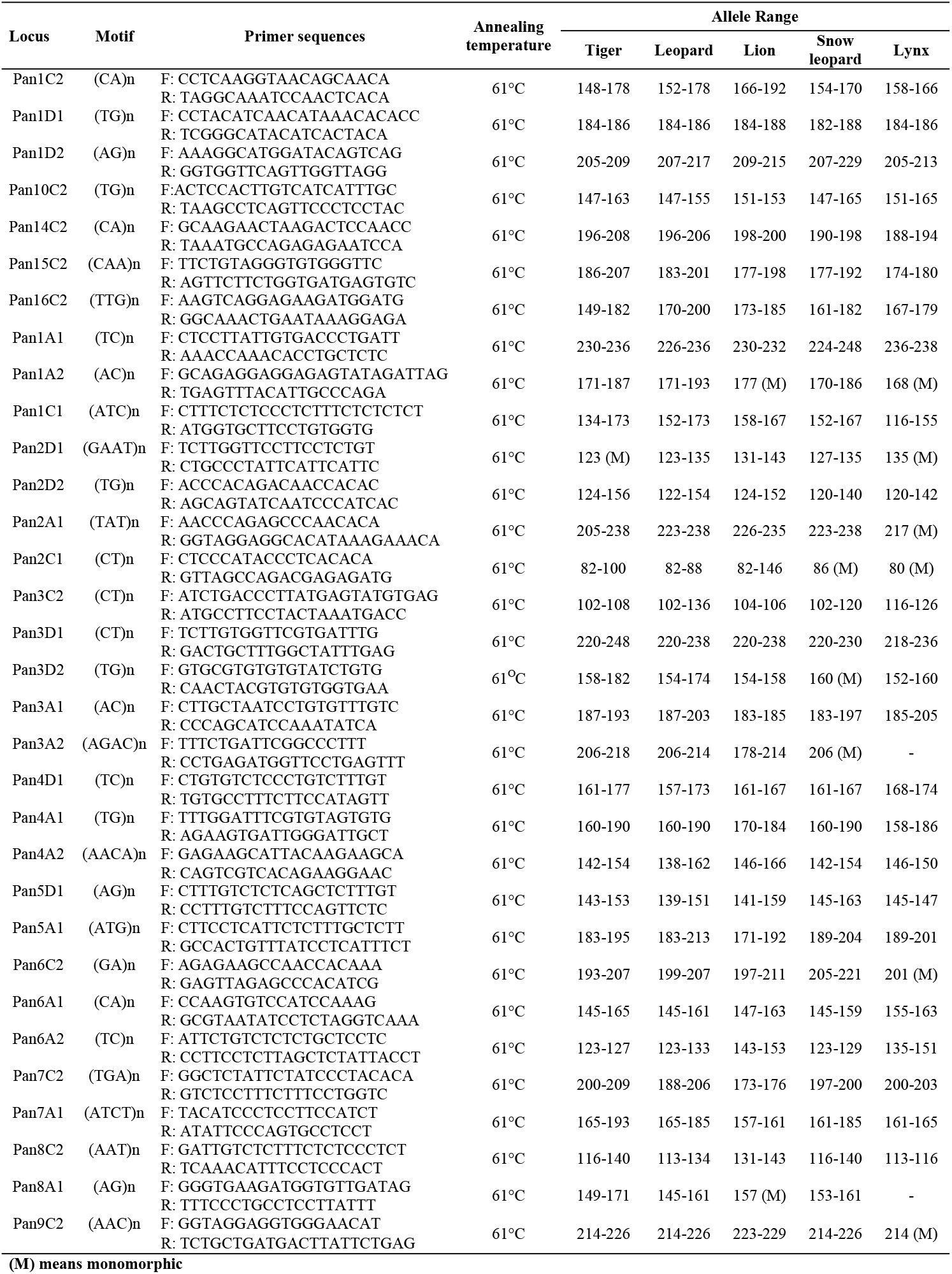
Description of 32 novel microsatellite loci developed for genus *Panthera*.

### Microsatellite polymorphism evaluation

The fluorescently labelled microsatellites were used to genotype 99 big cat individuals and 7 lynxes. Loci Pan3A2 and Pan8A1 failed to produce scorable profiles in lynx samples and thus, were assigned zero allelic value (Table 2). Overall, all loci were found to be polymorphic (4 to 18 alleles/locus), but some showed no variations within species - Pan2D1 in tiger and lynx; Pan1A2 and Pan8A1 in lion; Pan3A2, Pan3D2, and Pan2C1 in snow leopard; and Pan1A2, Pan2A1, Pan2C1, Pan6C2, and Pan9C2 in lynx (Table 2). The species wise microsatellite characteristics and polymorphism are as follows:

**Table 2:** Characterizationof 32 polymorphic microsatellite loci in four big cat species and Lynx.

| Locus | Number of alleles |  |  |  |  | Heterozygosity (observed) |  |  |  |  | Heterozygosity (expected) |  |  |  |  | Polymorphic information content |  |  |  |  |
| --- | --- | --- | --- | --- | --- | --- | --- | --- | --- | --- | --- | --- | --- | --- | --- | --- | --- | --- | --- | --- |
|  | Tiger | Leopard | Lion | Snow leopard | Lynx | Tiger | Leopard | Lion | Snow leopard | Lynx | Tiger | Leopard | Lion | Snow leopard | Lynx | Tiger | Leopard | Lion | Snow leopard | Lynx |
| Pan10C2 | 4 | 4 | 2 | 6 | 4 | 0.03 | 0.00 | 0.29 | 0.25 | 0.43 | 0.19 | 0.42 | 0.45 | 0.78 | 0.71 | 0.18 | 0.38 | 0.34 | 0.70 | 0.62 |
| Pan14C2 | 6 | 6 | 2 | 4 | 4 | 0.50 | 0.31 | 0.00 | 0.38 | 0.50 | 0.65 | 0.68 | 0.11 | 0.69 | 0.71 | 0.58 | 0.61 | 0.10 | 0.59 | 0.60 |
| Pan15C2 | 6 | 7 | 5 | 3 | 2 | 0.32 | 0.55 | 0.53 | 0.14 | 0.17 | 0.45 | 0.69 | 0.73 | 0.28 | 0.17 | 0.42 | 0.64 | 0.66 | 0.24 | 0.14 |
| Pan16C2 | 8 | 5 | 4 | 5 | 5 | 0.50 | 0.55 | 0.69 | 0.38 | 0.86 | 0.70 | 0.67 | 0.68 | 0.73 | 0.82 | 0.64 | 0.59 | 0.59 | 0.64 | 0.73 |
| Pan1A1 | 4 | 4 | 2 | 7 | 2 | 0.45 | 0.20 | 0.47 | 0.50 | 0.17 | 0.56 | 0.48 | 0.48 | 0.69 | 0.17 | 0.49 | 0.44 | 0.36 | 0.63 | 0.14 |
| Pan1A2 | 5 | 8 | 1 | 4 | 1 | 0.06 | 0.64 | 0.00 | 0.13 | 0.00 | 0.33 | 0.88 | 0.00 | 0.44 | 0.00 | 0.30 | 0.85 | 0.00 | 0.39 | 0.00 |
| Pan1C1 | 6 | 7 | 3 | 3 | 3 | 0.03 | 0.28 | 0.53 | 0.00 | 0.17 | 0.62 | 0.68 | 0.50 | 0.43 | 0.62 | 0.56 | 0.63 | 0.41 | 0.37 | 0.51 |
| Pan1C2 | 10 | 8 | 4 | 4 | 4 | 0.26 | 0.53 | 0.29 | 0.13 | 0.67 | 0.56 | 0.86 | 0.51 | 0.44 | 0.73 | 0.53 | 0.83 | 0.45 | 0.39 | 0.62 |
| Pan1D1 | 2 | 2 | 2 | 3 | 2 | 0.00 | 0.08 | 0.69 | 0.00 | 0.00 | 0.08 | 0.28 | 0.51 | 0.43 | 0.43 | 0.07 | 0.24 | 0.37 | 0.37 | 0.31 |
| Pan1D2 | 3 | 3 | 3 | 3 | 3 | 0.06 | 0.11 | 0.71 | 0.00 | 1.00 | 0.18 | 0.52 | 0.68 | 0.55 | 0.67 | 0.16 | 0.43 | 0.58 | 0.45 | 0.54 |
| Pan2A1 | 5 | 5 | 4 | 4 | 1 | 0.29 | 0.39 | 0.63 | 0.25 | 0.00 | 0.58 | 0.80 | 0.70 | 0.52 | 0.00 | 0.48 | 0.75 | 0.62 | 0.44 | 0.00 |
| Pan2C1 | 4 | 3 | 4 | 1 | 1 | 0.03 | 0.30 | 0.11 | 0.00 | 0.00 | 0.20 | 0.53 | 0.30 | 0.00 | 0.00 | 0.19 | 0.46 | 0.28 | 0.00 | 0.00 |
| Pan2D1 | 1 | 4 | 2 | 2 | 1 | 0.00 | 0.00 | 0.46 | 0.00 | 0.00 | 0.00 | 0.42 | 0.37 | 0.23 | 0.00 | 0.00 | 0.38 | 0.29 | 0.20 | 0.00 |
| Pan2D2 | 9 | 11 | 8 | 4 | 7 | 0.37 | 0.54 | 0.73 | 0.13 | 0.71 | 0.69 | 0.76 | 0.84 | 0.44 | 0.88 | 0.65 | 0.72 | 0.79 | 0.39 | 0.79 |
| Pan3A1 | 4 | 5 | 2 | 5 | 4 | 0.15 | 0.10 | 0.00 | 0.86 | 1.00 | 0.57 | 0.50 | 0.33 | 0.79 | 0.77 | 0.48 | 0.43 | 0.27 | 0.69 | 0.66 |
| Pan3A2 | 4 | 3 | 3 | 1 | 0 | 0.22 | 0.23 | 0.07 | 0.00 | 0.00 | 0.48 | 0.36 | 0.20 | 0.00 | 0.00 | 0.42 | 0.32 | 0.19 | 0.00 | 0.00 |
| Pan3C2 | 4 | 5 | 2 | 4 | 4 | 0.36 | 0.27 | 0.00 | 0.25 | 0.67 | 0.53 | 0.63 | 0.12 | 0.35 | 0.56 | 0.47 | 0.56 | 0.11 | 0.31 | 0.48 |
| Pan3D1 | 9 | 4 | 2 | 4 | 6 | 0.69 | 0.08 | 0.06 | 0.13 | 1.00 | 0.76 | 0.28 | 0.06 | 0.53 | 0.84 | 0.72 | 0.26 | 0.06 | 0.46 | 0.74 |
| Pan3D2 | 11 | 6 | 2 | 1 | 3 | 0.78 | 0.29 | 0.13 | 0.00 | 0.17 | 0.84 | 0.46 | 0.13 | 0.00 | 0.44 | 0.80 | 0.43 | 0.11 | 0.00 | 0.36 |
| Pan4A1 | 10 | 8 | 6 | 7 | 2 | 0.61 | 0.46 | 0.79 | 0.71 | 0.00 | 0.76 | 0.60 | 0.70 | 0.89 | 0.30 | 0.71 | 0.56 | 0.64 | 0.80 | 0.24 |
| Pan4A2 | 3 | 5 | 2 | 3 | 2 | 0.08 | 0.44 | 0.06 | 0.13 | 0.17 | 0.08 | 0.51 | 0.06 | 0.34 | 0.17 | 0.08 | 0.42 | 0.06 | 0.29 | 0.14 |
| Pan4D1 | 6 | 8 | 4 | 2 | 4 | 0.60 | 0.68 | 0.50 | 0.20 | 0.50 | 0.67 | 0.80 | 0.46 | 0.20 | 0.71 | 0.62 | 0.75 | 0.41 | 0.16 | 0.60 |
| Pan5A1 | 4 | 6 | 3 | 5 | 4 | 0.08 | 0.47 | 0.29 | 0.25 | 0.71 | 0.21 | 0.62 | 0.53 | 0.45 | 0.69 | 0.20 | 0.55 | 0.43 | 0.40 | 0.59 |
| Pan5D1 | 6 | 6 | 6 | 6 | 2 | 0.35 | 0.38 | 0.35 | 0.29 | 0.00 | 0.63 | 0.48 | 0.66 | 0.75 | 0.36 | 0.59 | 0.44 | 0.60 | 0.66 | 0.27 |
| Pan6A1 | 10 | 9 | 6 | 5 | 3 | 0.60 | 0.50 | 0.41 | 0.29 | 0.75 | 0.82 | 0.84 | 0.37 | 0.73 | 0.61 | 0.78 | 0.80 | 0.35 | 0.63 | 0.47 |
| Pan6A2 | 3 | 5 | 5 | 3 | 6 | 0.41 | 0.25 | 0.38 | 0.13 | 1.00 | 0.57 | 0.54 | 0.63 | 0.24 | 0.89 | 0.49 | 0.49 | 0.55 | 0.22 | 0.79 |
| Pan6C2 | 5 | 3 | 3 | 3 | 1 | 0.30 | 0.00 | 0.11 | 0.13 | 0.00 | 0.55 | 0.25 | 0.11 | 0.24 | 0.00 | 0.49 | 0.23 | 0.10 | 0.22 | 0.00 |
| Pan7A1 | 8 | 6 | 2 | 3 | 2 | 0.73 | 0.62 | 0.20 | 0.13 | 0.50 | 0.83 | 0.70 | 0.19 | 0.43 | 0.41 | 0.80 | 0.65 | 0.16 | 0.35 | 0.31 |
| Pan7C2 | 4 | 4 | 2 | 3 | 2 | 0.40 | 0.36 | 0.06 | 0.33 | 0.67 | 0.62 | 0.39 | 0.34 | 0.32 | 0.55 | 0.53 | 0.35 | 0.27 | 0.27 | 0.38 |
| Pan8A1 | 3 | 3 | 1 | 2 | 0 | 0.04 | 0.00 | 0.00 | 0.00 | 0.00 | 0.17 | 0.24 | 0.00 | 0.67 | 0.00 | 0.16 | 0.22 | 0.00 | 0.38 | 0.00 |
| Pan8C2 | 7 | 4 | 3 | 4 | 2 | 0.38 | 0.04 | 0.62 | 0.14 | 0.60 | 0.69 | 0.24 | 0.59 | 0.50 | 0.47 | 0.64 | 0.22 | 0.47 | 0.43 | 0.33 |
| Pan9C2 | 4 | 4 | 3 | 3 | 1 | 0.13 | 0.03 | 0.33 | 0.38 | 0.00 | 0.34 | 0.26 | 0.59 | 0.51 | 0.00 | 0.31 | 0.24 | 0.48 | 0.43 | 0.00 |

#### Tiger (*Pantheratigris*)

We genotyped 41 tiger individuals of wild and captive origin. They were collected from India (n = 8), Russia (n = 4), and South Korea (n = 29, zoo individuals). We pooled the samples from different locations to make a single tiger population to study polymorphism of the markers. Though not appropriate, this was the best possible way as (1) there were not enough Russian tiger samples to perform genetic analysis independently, (2) the ancestry of most of the zoo tigers was presumed to be hybrid (Bengal tiger and Amur tiger) ^48^, (3) Amur tiger and Bengal tiger are two ecotypes of a same subspecies ^49^. The number of alleles per locus ranged from 1 to 11 (mean: 5.6) with a mean expected heterozygosity of 0.5 (0.00 – 0.84). Twenty-six of 32 loci deviated significantly from HWE (Hardy-Weinberg Equilibrium) after Bonferroni correction (adjusted p-value < 0.002, Table S2), and null alleles were detected in 24 loci (threshold limit of 10%, Table S2). Deviation from HWE was expected due to Wahlund effect. Overall, the markers were found to be polymorphic (except Pan2D1) with a mean polymorphic information content (PIC) of 0.45. Fourteen markers were found to have PIC ≥ 0.5, indicating their informative nature and utility for conservation genetic studies (Table 2).

#### Leopard (*Pantherapardus*)

A total of 32 individuals belonging to the wild (India and Russia) and captivity (South Korea) were genotyped. Overall, markers were polymorphic in leopards with mean allelic diversity of 5.3 (2 – 11 alleles/locus) and average expected heterozygosity of 0.54 (0.24 – 0.88). Seven loci (Pan1A2, Pan1C1, Pan1C2, Pan1D2, Pan5D1, Pan6A1, and Pan6C2) in zoo leopards and one locus (Pan2D1) in Amur leopard sampled from Russia deviated significantly from HWE after Bonferroni correction (adjusted p-value < 0.002, Table S2). According to studbook records, all leopards sampled from Korean zoos belong to Indochinese subspecies (*Panthera pardus delacouri*). Thus, the probable HWE deviation may have resulted due to higher average relatedness or hybrid ancestry. Null alleles (≥10%) were detected in 19, 18, and 8 loci in leopards sampled from Russia, Korean zoos, and India (Table S2). However, there were inconsistencies in their occurrence in three tested populations. Thus, there is high probability of discovery of additional alleles in these developed markers, if tested with a greater number of samples. Fourteen of the 32 markers were found suitable for conservation genetic studies with PIC ≥ 0.5 (Table 2).

#### Lion (*Pantheraleo*)

A total of 18 captive African lions from Korean zoos were genotyped. Out of 32 loci, 2 were monomorphic and 30 were polymorphic loci, with the number of alleles ranging from 1 to 8 (mean = 3.2). The mean expected heterozygosity was 0.4 (0.00 - 0.84) for lions. We did not observe any significant deviation from HWE after Bonferroni correction (adjusted p-value < 0.002) in any loci (Table S2). Null alleles were detected in 9 loci (≥10%, Table S2). The mean polymorphic information content was estimated to 0.35, with 8 loci having PIC > 0.5 (Table 2).

#### Snow leopard (*Pantherauncia*)

Snow leopards (n = 8) were sampled from the wild (Mongolia) and zoo (Korea). All these samples were considered as a single population during genetic analysis as there were not enough samples from the wild or captivity to be considered as distinct populations. Moreover, Korean zoos sourced snow leopards from Mongolia.

In twenty-nine polymorphic microsatellites, the number of the alleles ranged from 2 to 7 (mean = 3.9), with mean expected heterozygosity of 0.5 (0.2 – 0.89). Locus Pan10C2 showed a significant deviation from HWE after Bonferroni correction (adjusted p-value < 0.002, Table S2). Null alleles were detected in 23 loci (≥10%, Table S2). The mean polymorphic information content was 0.4 with eight loci having PIC > 0.5 (Table 2).

#### Lynx (*Lynx lynx*)

Twenty-six loci were found polymorphic for Eurasian lynx with allele ranging from 2 to 7 (mean = 3.4) and mean expected heterozygosity being 0.57 (0.16 – 0.89). There was no sign of HWE deviation in tested loci after Bonferroni correction (adjusted p-value < 0.002, Table S2). Only 7 loci had null alleles above the threshold of 10% (Table S2). Twelve markers had PIC ≥ 0.5 (Table 2).

### Establishment of a universal microsatellite marker system for big cat species

This study aims to propose a universal microsatellite marker system capable of undertaking individual identification and geographic assignments of big cat seizures. We understand that the loci with higher expected heterozygosity (He) are more useful for individual identification. Similarly, loci with PIC values higher than 0.5 are considered informative enough for estimating genetic diversity. In our study, the locus wise heterozygosity and PIC varied across the species. We selected fifteen microsatellite loci based on the comparative marker’s PIC, heterozygosity, and allele diversity (Table 3). These loci showed no signs of linkage disequilibrium (LD) with big cats’ wild populations. The average PIC of 15 markers was 0.48, 0.50, 0.54, and 0.56 for the snow leopard, lion, tiger, and leopard, respectively. The cumulative power of discrimination among unrelated individuals (P_ID_) was found to be 5.2X10^-10^, 7.9X10^-10^, 3.0X10^-11^,and 5.2X10^-12^ for lion, snow leopard, tiger, and leopard, respectively, using the recommended panel of 15 microsatellites. Similarly, the cumulative power of discrimination among siblings (P_ID_ sib) was found to be 1.5X10^-4^, 7.8X10^-5^, 3.3X10^-5^, and 2.5X10^-5^ for the snow leopard, lion, tiger, and leopard, respectively.

**Table 3:** Probability of identity for unrelated samples (PID) and for full siblings (PID sib) in 15 microsatellite loci.

| **Locus** | **Tiger** |  | **Leopard** |  | **Lion** |  | **Snow leopard** |  |
| --- | --- | --- | --- | --- | --- | --- | --- | --- |
|  | P<sub>ID</sub> | P<sub>ID</sub> sib | P<sub>ID</sub> | P<sub>ID</sub> sib | P<sub>ID</sub> | P<sub>ID</sub> sib | P<sub>ID</sub> | P<sub>ID</sub> sib |
| Pan6A1 | 6.15E-02 | 3.62E-01 | 5.58E-02 | 3.54E-01 | 4.22E-01 | 6.76E-01 | 1.48E-01 | 4.50E-01 |
| Pan2D2 | 1.27E-01 | 4.43E-01 | 8.59E-02 | 4.00E-01 | 6.04E-02 | 3.60E-01 | 3.70E-01 | 6.36E-01 |
| Pan5A1 | 6.30E-01 | 8.02E-01 | 2.08E-01 | 4.98E-01 | 3.23E-01 | 5.76E-01 | 3.52E-01 | 6.27E-01 |
| Pan1C2 | 2.20E-01 | 5.31E-01 | 4.23E-02 | 3.38E-01 | 3.03E-01 | 5.79E-01 | 3.70E-01 | 6.36E-01 |
| Pan5D1 | 1.77E-01 | 4.85E-01 | 3.13E-01 | 5.92E-01 | 1.66E-01 | 4.73E-01 | 1.24E-01 | 4.34E-01 |
| Pan7C2 | 2.28E-01 | 5.02E-01 | 4.17E-01 | 6.64E-01 | 5.06E-01 | 7.13E-01 | 5.21E-01 | 7.34E-01 |
| Pan4A1 | 1.02E-01 | 4.01E-01 | 1.95E-01 | 5.02E-01 | 1.45E-01 | 4.48E-01 | 5.34E-02 | 3.50E-01 |
| Pan1C1 | 2.01E-01 | 4.94E-01 | 1.45E-01 | 4.55E-01 | 3.38E-01 | 5.95E-01 | 3.88E-01 | 6.44E-01 |
| Pan2A1 | 2.71E-01 | 5.33E-01 | 8.42E-02 | 3.81E-01 | 1.67E-01 | 4.51E-01 | 3.07E-01 | 5.85E-01 |
| Pan3A1 | 2.74E-01 | 5.39E-01 | 3.24E-01 | 5.85E-01 | 5.14E-01 | 7.18E-01 | 1.16E-01 | 4.12E-01 |
| Pan6A2 | 2.63E-01 | 5.32E-01 | 2.63E-01 | 5.49E-01 | 2.13E-01 | 4.98E-01 | 6.10E-01 | 7.89E-01 |
| Pan1A1 | 2.60E-01 | 5.38E-01 | 3.15E-01 | 5.92E-01 | 3.95E-01 | 6.16E-01 | 1.47E-01 | 4.63E-01 |
| Pan15C2 | 3.31E-01 | 6.13E-01 | 1.38E-01 | 4.48E-01 | 1.36E-01 | 4.30E-01 | 5.70E-01 | 7.65E-01 |
| Pan16C2 | 1.45E-01 | 4.44E-01 | 1.79E-01 | 4.69E-01 | 1.84E-01 | 4.69E-01 | 1.41E-01 | 4.45E-01 |
| Pan8C2 | 1.39E-01 | 4.48E-01 | 5.98E-01 | 7.82E-01 | 2.82E-01 | 5.37E-01 | 3.24E-01 | 6.02E-01 |
| **Cumulative** | **3.03E-11** | **3.30E-05** | **5.17E-12** | **2.54E-05** | **5.21E-10** | **7.82E-05** | **7.92E-10** | **1.48E-04** |

## Discussion

Even with the development of more sophisticated and elaborate markers such as SNPs, microsatellites are still considered the best tool to study conservation genetics due to their codominant inheritance pattern and hypervariability. There are two kinds of microsatellites – species-specific and heterologous. The former is developed for a species of interest, while the latter is screened from a pool of STR loci that were previously described for other species. Geneticists have used both species-specific and heterologous microsatellites to study the genetic diversity and population structures of big cats^31,32,50–52^. However, the use of heterologous markers is more prevalent due to the availability of a limited number of species-specific STRs. Mishra et al. (2014) compared the polymorphism of species-specific vs. cross-specific markers in Bengal tiger and concluded the former’s superiority over the latter ^53^. Moreover, the chances of genotyping errors due to mispriming, false alleles, and null alleles are lesser with species-specific STRs. In this study, the genome sequences of seven big cat individuals belonging to four species were analysed rapidly to identify and develop thirty-two polymorphic loci. The procedure of microsatellite development involved four steps: (1) mapping of big cat genomes on the assembled reference genome of the domestic cat to develop a multiple sample construct, (2) screening of the unique variant sites from the multiple sample construct, (3) scanning of unique variants to identify the polymorphic STR loci with conserved flanking regions, and (4) designing of PCR primers for these loci and evaluation of polymorphism with the collected samples (Figure 1). Since the whole process involved comparative genome analysis and selection of universally located STRs with conserved flanking regions, the developed microsatellite markers were regarded as species-specific for all the four target big cat species. This makes our study a pioneering attempt to develop microsatellite markers for a genus. The autosomal location of each marker was assigned based on the karyotype of the domestic cat as its karyotype is reported to be similar to that of *Panthera* species. The microsatellite markers were named according to the genus *Panthera* (Pan) and autosome location (A1, A2, D1, etc., Table 1). For example, Pan10C2, Pan14C2, Pan15C2, and Pan16C2 are markers located on chromosome C2 in all Panthera species. Microsatellites were found to be located on six of the eighteen autosomal chromosomes, thereby ensuring at least 33% genome coverage.

We developed fluorescently labelled primer pairs for 32 novel microsatellite loci. Their polymorphism potential was evaluated with the DNA samples of four big cat species and lynx. All markers amplified successfully and produced scorable profiles with tiger, lion, leopard, and snow leopard. However, the profiles of Pan3A2 and Pan8A1 were un-scorable with lynx samples. The faulty genotyping profiles could have resulted due to non-target primer annealing. The increased phylogenetic distance between the source (big cats: genus *Panthera*) and target (Eurasian lynx: genus *Lynx*) species greatly reduces transferability of markers ^54^.

All markers were found polymorphic in leopards. Pan2D2 in tiger, Pan1A2, and Pan8A1 in lion and Pan3A2, Pan3D2, and Pan8A1 in snow leopard were monomorphic. Mean allelic diversity was found highest for tigers followed by leopard, lion, and lynx (Table 2). The evidence of null alleles in several locus suggests that more alleles may be discovered. We also reported significant deviation from HWE in several loci in tiger, leopard, and snow leopard (Table S2). This could have resulted due to pooling of samples of different subspecies or populations into one group (Wahlund effect) or the analysis of first-degree relatives. Both are possible in our case as we sampled captive individuals and pooled samples based on broad geographical limits. Therefore, we recommend further evaluation of these novel markers with more samples before drawing a conclusion about their polymorphism potential.

Microsatellite polymorphism levels vary greatly across populations and species. Markers with PIC greater or around 0.5 were considered suitable for genetic studies. Fourteen each in tiger and leopard, and 8 each in lion and snow leopard had PIC values greater than the threshold (Table 2).

Identification of affected species, the responsible perpetrators, and their methods of killing are important aspects of wildlife forensic investigations. However, wildlife managers are only interested in the information about the affected species and population (source). Knowledge of the origin of the confiscated wildlife helps in the initiation of remedial actions in a timely manner. Microsatellite markers are great tools for the scientists and technicians involved in the investigation of wildlife poaching and trade cases. Microsatellite-based genetic IDs are useful to ascertain the number of affected (killed) individuals. The same information can then be used to reveal the source population (geographic assignment).

Tigers are the most illegally traded big cat species. In the past few decades, the increasing substitution of tiger parts with that of other big cat species has been observed. Except for pelt, commercially traded parts of big cats such as claw, bone, whisker, meat, canine, etc. are morphologically indistinguishable at the species level. In 2015, Mondol et al. successfully demonstrated the use of microsatellite markers to infer the source of origin of the leopard seizures from India ^42^. Similarly, Zou et al. (2015) proposed a panel of microsatellites for tigers to identify individuals and subspecies ^43^. In both studies, researchers generated a microsatellite-based genetic signature of all candidate populations (or subspecies) on their own, as the available information in the published domain was incompatible due to the use of different STR loci. Thus, to ensure the adoption of the microsatellite-based approach in forensic investigations, there is a need for the use of a unified DNA typing methodology for individual identification and establishment of genetic signatures. Moreover, the use of an established and universal methodology is more convincing during court proceedings. Here, we proposed a universal microsatellite panel for four big cat species that are most affected by illegal trade and are often traded with the same covert identity. The panel includes 15 microsatellite loci, distributed over six chromosomes, and providing approximately 33% genome coverage (Table 3). Cumulatively, these markers have a high discriminatory power of one in a million for unrelated individuals and one in a thousand for siblings (Table 3). Such a high degree of discriminatory power also makes this panel suitable for population genetic studies. In the wild, more than two big cat species often inhabit the same region or country simultaneously (e.g., tiger, leopard, lion, and snow leopard in India; lion and leopard in Africa; tiger, leopard, and snow leopard in Russia). The universal marker system for all the big cat species will reduce the necessary reagent cost and technical burden of researchers working on different big cat species in a laboratory or a network of laboratories. This will also promote data exchange and cooperative research. The similar range of annealing temperatures of primers (Table 1) for the markers in this study will be useful for developing a multiplex PCR system. Besides, since the markers are developed by mining the polymorphic STR loci with conserved flanking regions using the assembled genomic sequence of the domestic cat as the reference sequence, most of the markers have the potential to be applied to a variety of other endangered cat species. The potential is exemplified by the use of lynx in this study; 30 out of 32 markers were successfully amplified using lynx samples. Hence, the proposed microsatellite panel is of great utility in establishing DNA fingerprints, population signatures, and wildlife forensics.

## Materials and methods

### Sample collection and DNA preparation

We collected biological samples of tiger, leopard, lion, and snow leopard from nature reserves, zoos, and sample repositories of India, Mongolia, Russia, and South Korea (Table S1). These include blood, muscle, faeces, shed hair, and DNA extracts. In our study, we also included DNA extracts of seven Mongolian Eurasian lynx (*Lynx lynx*) to assess the utility of the developed markers over other cat species (Table S1). This experimental work was conducted with permission by the Conservation Genome Resource Bank for Korean Wildlife (CGRB)that provided the biological samples of wild cats for this study. All samples were legally and ethically collected and wherever applicable, the necessary permissions and permits were obtained from competent authorities. The procedures involving animal samples followed the guidelines by Seoul National University Institutional Animal Care and Use Committee (SNU IACUC).The species identity of each of the sourced samples were reverified using conservation genetic tools i.e., amplifying either species-specific primers ^55,56^ or by sequence analysis of Cyt b gene using universal primers ^57^.

The commercial column-based DNA extraction kits were employed to extract DNA following the recommended protocols. The whole process was carried out in a sterile environment of a dedicated laboratory to avoid any chance of contamination. Further, a positive and a negative control per experimental setup were included. Post extraction, DNA was resolved on 0.8% agarose gel to assess quality and quantity. Finally, the DNA was preserved at −20°Cfor long term storage.

### Microsatellite development for genus Panthera

In our study, we analysed previously published genome sequences of seven big cat individuals ^58,59^. These include three tigers, two lions, a leopard, and a snow leopard. Additionally, we downloaded the assembled genome of domestic cat, Felcat6.2 ^60^, that served as areference. The whole process has been schematised in Figure 1.

Each genome was processed independently for the variant calling. The FASTQ reads of the individual genome were mapped on the assembled reference genome (Felcat6.2) with the BWA-MEM ^61^ using the default options. Duplicates were marked using Picard Tools. Thereafter, the variant sites were assessed using the Samtools mpileup ^62^ and consensus sequences were generated for each species. A multiple sample construct was developed to make the genomes of different species comparable and to identify the variable sites. Samples without variants at the position were assigned the reference allele with the related coverage from the sample. The variants were then filtered based upon the following criteria: no heterozygous status for any sample, depth greater than or equal to 4 for all samples at that position (DP>4), and the number of different alleles among all the samples present should be greater than a specified value (like 3, 4, 5, or 6 unique alleles) out of the possible total. The resulting variants were considered as the potential target variants. These were then parsed for unique sites since it is possible to have variants called from different samples at the same site. The unique target variant sites were then expanded to +150bp around the sites to create 301bp regions for downstream primer design. The nucleotide sequence of the Felis catus reference at those covered regions was extracted by BEDTools ^63^, and variant sites were replaced with the longest allele from all possible alleles at the site.

The program MSDB ^46^ was used to screen the perfect STR repeats of 1-6 bp having a minimum repeat number of 12, 7, 5, 4, 4, and 4 for mono-, di-, tri-, tetra-, penta-, and hexanucleotide microsatellites respectively, from the unique target variant sequences. The repeats were classified into classes based on their start position and reverse complements. For example, TGG contains TGG, GGT, GTG, ACC, CCA, and CAC in different reading frames or on complementary strands. Microsatellite average length, total counts, frequency (loci/Mb), and density (loci/bp) of the motif were analysed ^46^. The sequences of microsatellite repeat regions that passed the selection criteria were used to design the primer sets using software Batch Primer 3 ^47^. The loci with long enough flanking regions (i.e., more than 20 bp) and with no single copy sequences were shortlisted for primer design. Further scanning was done using Clustal X1.83 ^64^ to ensure that the microsatellite should not be published earlier. The criteria for searching of the primers were as follows: (1) PCR product should range from 80 to 250 base pair considering the utility of developed markers with samples yielding low quality DNA, (2) primers melting temperature (Tm) should range from 52°C to 62°C (optimal 55°C), (3) primer GC content should range from 40% to 60%, and (4) number of returns i.e. number of primer pairs generated for each unique target variant sequence should be four. The rest of the parameters were set to default.

Non-labelled primer pairs were synthesised for loci qualifying the primer designing and selection criteria. These primers were subsequently tested for PCR amplification with one sample each, of tiger, leopard, lion, and snow leopard. Gradient PCR (annealing temperature, T_a_ - 52°C to 62°C, reaction volume – 10 μL and primer concentration – 5 pm each) was performed independently for each primer pair. Primer pairs producing a single product band of expected size during PCR amplification were shortlisted for fluorescent dye labelling (forward primers) with one of four fluorescent dyes (6-FAM, VIC, NED, or PET, Invitrogen, South Korea) to perform fragment analysis using Applied Biosystems 3130 Genetic Analyser. During primer dye-labelling, due consideration was given to avoid dye range and product size overlap.

### Microsatellite polymorphism evaluation

Fluorescently labelled microsatellites were tested for their polymorphism potential in an independent PCR assay with 106 samples of big cats and lynx. In a reaction, the total volume was 10 μl, with 30-35 ng of extracted DNA, 1X PCR buffer, 0.25 mM dNTP mix, 0.5 U of i- StarTaq™ DNA polymerase (iNtRON Biotechnology, Inc), and 0.4 μM of each forward and reverse primer. The thermal profile of the amplification was as follows: initial denaturation at 94°C for 2 minutes, followed by 40 cycles of denaturation at 94°C for 40 seconds, annealing at 61°C for 40 seconds, extension at 72°C for 45 seconds, with one cycle of final extension for 30 minutes at 72°C. The amplified PCR products were checked on 2% agarose, diluted (1:20) with distilled water, pooled based on dye label and product size, and subjected to fragment analysis with an Applied Biosystems 3130 Genetic Analyzer. The alleles were scored with Gene Mapper 3.7 (Applied Biosystems).

During analysis, the samples were classified into sets: (1) based on species - 5 populations, and (2) based on species and geographic origin - 10 populations (Table S1). The microsatellite data was analysed for possible genotyping errors of scoring and stuttering with MicroChecker 2.2.3 ^65^. Conformance with HWE and level of LD were assessed using Genepop 1.2 ^66^. The p-values for HWE and LD were corrected for multiple comparisons by applying a sequential Bonferroni correction ^67^. Null allele frequencies were determined with the Dempsters EM method implemented in Genepop 1.2 ^66^. The software CERVUS was used to calculate the locus wise observed and expected frequency of alleles and heterozygosity, and the PIC for each population ^68,69^. Allele range was calculated for each of the markers by compiling the observed allele range of all species. Program Gimlet 1.3.3 was used to estimate P_ID_ for unrelated samples and more conservative P_ID_ sib to test the discriminatory power of sets with a different number of markers.

## Acknowledgements

We extend our thanks to administrative heads of Research Institute for Veterinary Science, Seoul National University College of Veterinary Medicine (Republic of Korea), Wildlife Institute of India (India), Land of the Leopard National Park (Russia), Seoul Grand Park Zoo (Republic of Korea), Everland Park (Republic of Korea), National University of Mongolia (Mongolia) for providing necessary permissions, samples, and facilities to carry out research. We are also thankful to Hanchan Park, Sujeet Singh, and Yunsun Lee for assistance in laboratory work. This work was supported by Brain Fusion Program of Seoul National University (No. 550-20140052), Bio Bridge Initiative grant of Ministry of Environment (2018-2019, Republic of Korea and Convention on Biological Diversity) and Indo-Korean Research Internship Program (2015) of National Research Foundation (Republic of Korea) and Department of Science and Technology (India).

## Author Contributions

HL, PP and KSK conceived and designed the experiments, AC, JB and PP did genome analysis, PP, JYH, DB and DJ performed experiments, HL, MHY, HKS, DB, JB, KSK, RS, MSM and SPG contributed reagents, materials, and analysis tools, and PP and JYH wrote manuscript with the help of other authors.

