## Supplementary material for "Whole genome survey of big cats (Genus: *Panthera*) identifies novel microsatellites of utility in conservation genetic study": Supplimentary Information

**Table S1:** Details of the samples collected for genetic analysis. All the samples were legally and ethically collected from zoos, national parks, wildlife institutions with permit from relevant authorities.

| | | Geographical location | | | |
|---|---|---|---|---|---|
| **Species** | Total | India | Republic of Korea* | Russia | Mongolia |
| **Tiger** | 41 | 8 | 29 | 4 | |
| **Leopard** | 32 | 8 | 16 | 8 | |
| **Lion** | 18 | | 18 | | |
| **Snow leopard** | 8 | | 2 | | 6 |
| **Lynx** | 7 | | | | 7 |

*from zoos

**Table S2:** Frequency (%) of null alleles across 32 novel microsatellite markers.

| Locus | Tiger | | | | Leopard | | | | Snow leopard | Lion | Lynx |
|---|---|---|---|---|---|---|---|---|---|---|---|
| | Overall | India | Russia | Zoo | Overall | India | Russia | Zoo | | | |
| Pan10C2 | 21.2* | - | 0.0 | - * | 35.5* | - | 58.5 | - | 42.2* | - | 13.0 |
| Pan14C2 | 9.9 | - | 22.2 | 8.7 | 24.6* | 63.7 | - | 15.1 | 35.1 | - | 0.0 |
| Pan15C2 | 10.1 | - | 31.5 | 0.0 | 9.1* | 0.0 | - | 7.3 | 23.6 | 7.4 | - |
| Pan16C2 | 16.4* | 16.7 | 31.5 | 12.4 | 14.5 | - | 30.2 | 42.8 | 17.8 | 3.1 | 0.0 |
| Pan1A1 | 0.0* | - | 40.0 | 65.4 | 23.0 | - | 85.1 | - | 14.6 | - | - |
| Pan1A2 | 26.0* | - | 40.0 | 0.0* | 16.5* | 27.5 | 18.9 | 14.4* | 31.2 | - | - |
| Pan1C1 | 38.6* | 52.5* | - | - * | 26.8* | 14.3 | 65.5 | 79.0* | 41.3 | 0.0 | 47.0 |
| Pan1C2 | 22.2* | 44.5* | - | 10.5 | 25.3* | 3.4 | 47.0 | 36.3* | 31.2 | 19.2 | 13.7 |
| Pan1D1 | - | - | - | - | - * | - | - | - | 87.1 | - | - |
| Pan1D2 | 93.6* | - | - | 94.4 | 31.5* | - | - | 26.3* | 47.0 | 22.9 | 31.5 |
| Pan2A1 | 21.6* | - | - | 16.9* | 33.7* | 38.2 | - | 7.3 | 36.8 | 6.0 | - |
| Pan2C1 | 21.8* | - | - | 17.6* | 39.1* | - | - | 47.9 | - | 24.5 | - |
| Pan2D1 | - | - | - | - | 87.1* | - | 67.8* | - | - | - | - |
| Pan2D2 | 25.5* | 0.0 | - | 25.8* | 14.9* | 24.3 | 21.5 | 0.0 | 31.2 | 20.0 | 15.0 |
| Pan3A1 | 29.7* | 29.0 | - | 48.7* | 28.7* | - | 20.7 | 23.6 | 28.6 | - | 0.0 |
| Pan3A2 | 19.7* | 25.1 | - | 25.2* | 19.2 | - | - | 2.4 | - | 17.6 | - |
| Pan3C2 | 16.9* | 7.9 | - | 22.1* | 24.3* | - | 26.9 | 33.3 | 18.3 | - | 0.0 |
| Pan3D1 | 8.8* | 20.7 | - | 7.4* | 29.6* | - | 50.8 | 22.4 | 31.2 | - | 0.0 |
| Pan3D2 | 9.7* | 0.0 | - | 16.0* | 20.2* | - | 41.3 | 26.8 | - | - | 0.0 |
| Pan4A1 | 11.8* | 0.0 | - | 12.2* | 11.7 | 19.8 | 20.7 | 10.9 | 15.9 | 0.0 | - |
| Pan4A2 | 0.0 | - | - | 0.0 | 4.0 | - | - | 0.0 | 0.0 | - | - |
| Pan4D1 | 11.9* | 19.5 | - | 21.8 | 6.4* | 0.0 | 47.0 | 0.0 | - | 0.0 | 17.6 |
| Pan5A1 | 20.6* | - | - | 17.6* | 8.9 | 0.0 | 25.1 | 12.8 | 20.7 | 44.6 | 18.9 |
| Pan5D1 | 25.3* | 14.5 | - | 16.1* | 13.3* | - | 26.8 | 0.0* | 33.0 | 20.6 | - |
| Pan6A1 | 23.4* | 0.0 | - | 27.7* | 19.2* | 0.0 | 14.6 | 25.9* | 44.3 | 0.0 | 0.0 |
| Pan6A2 | 12.8 | 0.0 | - | 9.6 | 21.3* | 31.5 | 16.8 | - | 0.0 | 20.2 | 8.8 |
| Pan6C2 | 26.0* | - | - | 26.0* | 34.4* | - | - | 38.7* | 22.4 | 0.0 | - |
| Pan7A1 | 7.0* | 0.0 | - | 3.9 | 18.0 | 16.7 | 0.0 | 27.5 | 29.0 | - | - |
| Pan7C2 | 18.0* | 62.3 | - | 19.2* | 9.8 | - | 4.7 | 10.0 | 0.0 | - | - |
| Pan8A1 | 17.9* | - | - | 19.5* | 93.4* | - | - | - | - | - | - |
| Pan8C2 | 20.9* | 7.6 | - | 16.7* | 23.7* | - | - | 19.5 | 32.8 | 0.0 | - |
| Pan9C2 | 24.6* | 22.4 | - | 26.0* | 92.0* | - | 87.1 | - | 72.3 | 24.1 | - |

**‘ - ’ – Insufficient alleles to calculate null allele frequency, * HWE deviation**
